# Inducible gene switches with memory in human T cells for cellular immunotherapy

**DOI:** 10.1101/346783

**Authors:** Deboki Chakravarti, Leidy D Caraballo, Benjamin H. Weinberg, Wilson W. Wong

## Abstract

Cell-based therapies that employ engineered T cells—including the expression of chimeric antigen receptors (CARs)—to target cancer cells have demonstrated promising responses in clinical trials. However, engineered T cell responses must be regulated to prevent severe side effects such as cytokine storms and off-target responses. Here we present a class of recombinase-based gene circuits that will enable inducible switching between two states of adoptive T cell therapy using an FDA-approved drug, creating a generalizable platform that can be used to control when and how strongly a gene is expressed. These circuits exhibit memory such that induced T cells will maintain any changes made even when the drug inducer is removed. This memory feature avoids prolonged drug inducer exposure, thus reducing the complexity and potential side effect associated with the drug inducer. We have utilized these circuits to control the expression of an anti-Her2-CAR, demonstrating the ability of these circuits to regulate CAR expression and T cell activity. We envision this platform can be extended to regulate other genes in T cell behavior for various adoptive T cell therapies.

## Introduction

T cells have emerged as a promising candidate for cell-based therapies, and engineering of T cells to express antigen-specific chimeric antigen receptors (CARs) have enabled programmable targeting of cancer cells^1-4^. Multiple clinical trials with CARs against B cell cancers have driven up to a 90% complete response rate in patients^5-8^, and several CAR T cell products targeting acute lymphoblastic leukemia (ALL) and non-Hodgkin lymphoma have been approved for clinical use in the United States.

Despite these promising clinical results, there are significant safety and efficacy considerations with CAR T cell therapies, often mirroring fundamental regulatory challenges of the immune system^9, 10^. For example, the immune system naturally seeks to prevent autoimmune reactions by selecting against highly autoreactive T cells during their development in the thymus^11^. However, engineered cancer-specific receptors often target markers that while overexpressed on tumor cells, may still be found at lower levels in healthy tissues^12^. These modified cells have the potential for an autoreactive “on-target, off-tumor” response, which has been observed and proven fatal in at least one clinical trial^13^. In addition, there are numerous regulatory checks that prevent the immune system from responding too strongly against pathogens and causing systemic harm, checks that may become disrupted by engineered T cells. CARs, in particular, can instigate a strong cytokine release in response to antigen stimulation, accelerating the immune response to potentially fatal levels. This cytokine release syndrome (CRS) has been observed in several CAR T cell clinical trials^7, 8^, and a regimen of immunosuppressive drugs is often required to ameliorate the response^5^.

These safety considerations point to the dangerous aspects of what is otherwise the major advantage of cell-based therapies: the ability to drive strong responses based on the cell’s own machinery. Targeted cytotoxicity is fundamental to the power of T cell therapies, but it is also the basis of its potential harms. This challenge is compounded by the immense cost—both in time and money—of cell-based therapies, which makes it difficult to iterate this therapy over and over again until it meets a patient’s individual needs.

T cell therapies would thus greatly benefit from the development of tools that allow greater control over cell behavior, enabling an already personalized therapy to be customizable towards a patient’s immediate and changing needs. Advances in genetic engineering and synthetic biology have provided significant insight into both the design and implementation of such controls in the form of synthetic receptors, protein-based switches, genetic circuits, and genome editing tools^9^. These tools have in turn been used to create “ON-Switch” CARs^14^, combinatorial activation systems^15,16^, doxycycline-inducible CARs^17^, antibody-inducible CARs^18^, kill switches^19-21^, pause switches^22^, tunable receptor systems^23-25^, proliferation switches^26^, and a universal “off the shelf” T cell^27^. These systems all reflect the tremendous potential of synthetic biology approaches in developing safer and more powerful forms of T cell therapy that can be customized to fit each patient.

While these technologies offer important forms of control over T cell behavior, their designs are accompanied by limitations. For example, while kill switches provide vital control in cases where the T cells are toxic to the patient, completely shutting off the therapy may be an extreme response for patients who only require a slight modification of the therapy to abrogate negative reactions. Some approaches are also only limited to certain types of therapy, such as the many CAR receptor designs that strictly provide added flexibility and control to CAR T cell therapy. In addition, certain inducible systems like the “ON Switch” CAR require the drug-inducer to be continuously provided to maintain the ON state, which may be less ideal if permanent changes are required for a patient. Furthermore, prolonged drug-inducer exposure may be detrimental to patients if the inducer has a less than ideal safety profile, even when the drug is FDA-approved. While these approaches will advance the scope of potential T cell therapies, developing further technologies that are compatible with them may help to expand their use.

To address the need for advanced control of T cell responses for various immunotherapy applications, we have developed a drug-inducible genetic circuit platform that is designed to be lentivirus-compatible, which will facilitate genetic modification of human primary T cells. Our system also has memory capability that reduces the need for prolonged drug administration to maintain gene expression level. We have adapted these circuits to control CAR expression, creating a variety of circuits, including an On Switch (ON), Off Switch (OFF), and Expression Level Switch (EXP) that controls CAR expression, alters T cell behavior, retains memory, and exhibits activity that can be tuned via drug dosage and duration. These gene switches represent the most versatile switches in T cells and have the potential to improve the safety and efficacy of T cell immunotherapy.

## Results

### Recombinase-based gene switch for controlling CAR expression

To implement a lentivirus-compatible, two-state switch with memory in T cells, we have adapted the recombinase-based flip-excision (FLEx) stable inversion switch for T cells. Recombinases are enzymes that can perform inversion or excision steps on DNA based on the relative orientation of DNA recognition sites. Recombinases were chosen for this work because they have demonstrated exceptional versatility and performance in engineered for gene regulation systems in mammalian cells^28^. The FLEx switch wasinitially designed using the Cre/*lox* system to regulate gene expression in mammalian cells via retroviral transduction of the switch^29^. This system relies upon the availability of orthogonal *lox* variant sites that are recognized by the Cre recombinase but do not interact with other variant sequences. Activation of the FLEx switch with recombinase begins with an unstable inversion step followed by a stable excision step, effectively removing one sequence of DNA and inverting another (**Fig. 1**). The overall product is a stable inversion switch that—when genes are encoded between the recombination sites—can stably alter gene expression via recombinase activity.

**Figure 1.**
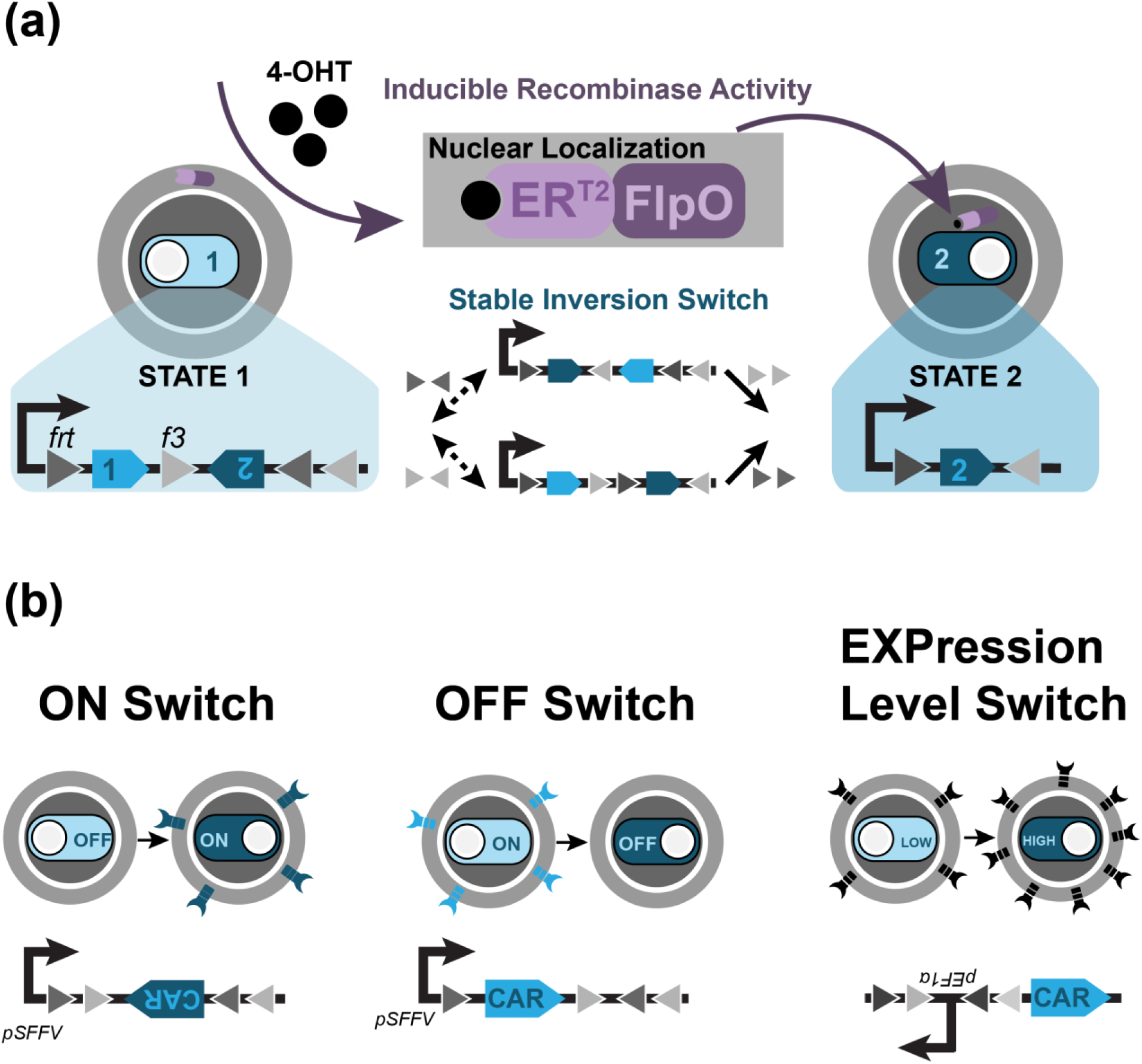
FlpO recombinase based FLEx switch design. (**a**) Mechanism of the 4-OHT-inducible FLEx switch. Induction with 4-OHT drives nuclear localization of the FlpO recombinase, initiating inversion and then excision upon the *frt/f3* recognition sites. By encoding genes representing State 1 and State 2 between the recognition sites, induction of FlpO activity stably shifts the cell from State 1 to State 2. (**b**) Design of the ON, OFF, and the EXPression level switch to control expression of CAR. The ON and OFF Switch express the CAR gene under State 1 and State 2 respectively. The EXP switch alters the orientation of the EF1α promoter relative to a CAR gene.

The FLEx switch exhibits several features that make it both applicable and beneficial towards T cell therapies. The stable inversion capability means that unlike a transcriptionally-inducible gene system, this circuit contains memory: when recombinase activity is terminated, changes made to the cells are maintained. This property is ideal for therapeutic strategies that seek a permanent change to T cell behavior without requiring continuous drug intake. It also enables changes to remain robust in response to rapid changes in proliferation that may dilute protein levels. In addition, the FLEx switch eschews the use of virus-incompatible genetic elements such as transcription termination sites that, while useful building blocks in complex recombinase-based logic systems ^28^, interfere with the reverse transcription process of viral integration and are incompatible with clinical settings that require viral transduction of T cells. The FLEx switch does not require transcriptional stop sites and is compatible with viral and transposon integration methods.

While the FLEx switch has been designed with the Cre/*lox* system, Cre exhibits toxicity^30, 31^ in mammalian cells due to the presence of pseudo-*loxP* sites in the genome. This genotoxicity requires careful tuning to be mitigated^32^. We initially developed a Cre/*lox*-based FLEx switch into Jurkat T cells and observed high toxicity upon Cre induction (**Supplementary Fig. 1**). We adapted the FLEx switch to operate with the FlpO/*frt* recombinase system instead, as the Flp recombinase has not been reported to be toxic^33, 34^, using the *frt* site and its variant *f3* site. A similar FlpO-based FLEx switch has been designed using other variant *frt* sites^35^. To control recombination, we used FlpO conjugated to the mutated estrogen receptor ER^T2 36^, which localizes the recombinase to the cytoplasm^29^. Upon induction with the drug 4-hydroxytamoxifen (4-OHT), the recombinase is localized to the nucleus, where it can act upon the FLEx switch (**Fig. 1a**). We chose the tamoxifen-inducible Flp for our circuit design because tamoxifen is an FDA-approved drug, which will facilitate implementation into the clinic. We have adapted the FlpOER^T2^/*frt* FLEx switch design to alter the orientation of CAR genes to create a stable ON and stable OFF switch (**Fig. 1b**). In addition, we have designed an “Expression Level” (EXP) switch that affects the orientation of the human EF1α promoter relative to the CAR gene, stably taking cells from low expression to high expression of CAR (**Fig. 1b**).

### Induction of recombinase activity drives changes in CAR expression

We transduced human primary CD4+ T cells with two viruses: one that contained a constitutively expressed FlpOER^T2^, and another expressing either the ON, OFF, or EXP switch controlling the expression of an αHer2-CAR. We sorted cells for FlpOER^T2^ expression and then induced with 4-OHT, a metabolite of tamoxifen, and observed changes in CAR expression via flow cytometry. All switches contained a CAR expressing an extracellular myc epitope tag that could be detected via antibody staining, but CAR expression in the EXP Switch was measured through an mCherry fluorescent tag directly conjugated to the CAR, which enabled rapid measurement. However, to reduce the potential of altered CAR degradation rates due to the mCherry fluorescent tag, ON and OFF Switch cells expressed a CAR that lacked the fluorescent tag, and instead CAR expression in these cells was measured by staining for the myc epitope.

All three circuits exhibited changes in anti-Her2 CAR expression in recombinase-positive CD4+ T cells within one day of induction (**Fig. 2**). The ON switch, in particular, demonstrated fast kinetics, reaching its maximal percentage of switched cells within one day. Meanwhile, OFF Switch cells demonstrated a loss in CAR+ cells within one day of induction, but the population required six days to stabilize (<10% cells expressing CAR). The slower dynamics of the OFF Switch compared to the ON Switch is likely due to the need to degrade and dilute CAR expression, making the OFF switch more reliant upon both growth and protein degradation rates.

**Figure 2.**
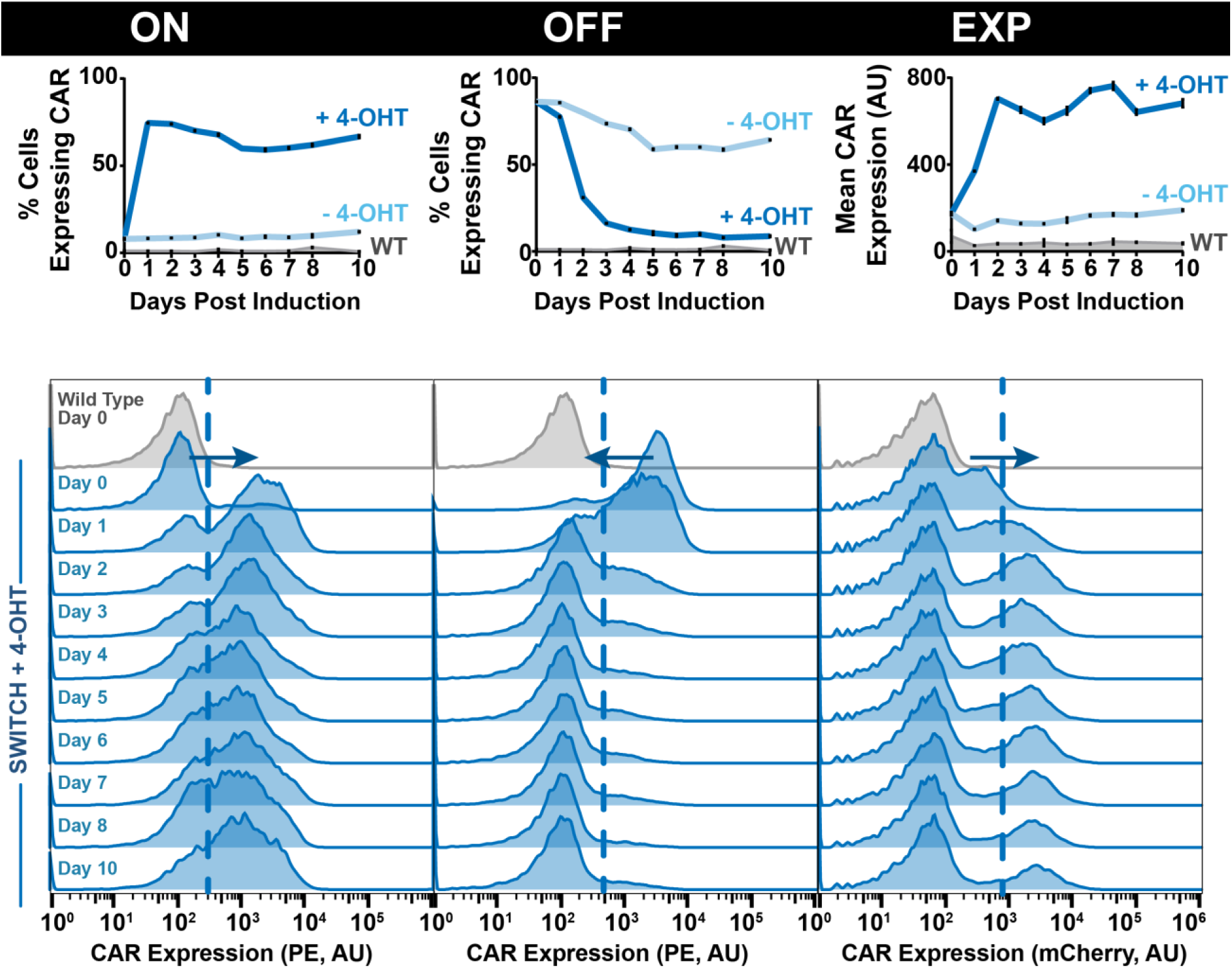
FlpO can be used to create ON, OFF, and EXPression level switches to control αHer2-CAR expression in human primary CD4+ T cells. *Top Panel*. Time course data for the recombinase switches with or without drug addition (4-OHT). Samples were obtained in triplicate from each induced or non-induced culture and then plotted as mean and standard deviation. The ON and OFF switches are presented as percent cell expressing the aHer2-CAR. The EXP switch is presented as the mean CAR expression level in arbitrary units (AU). *Bottom Panel*. The histogram of CAR expression level (AU) at different days after 4-OHT addition. Dashed vertical line illustrates the CAR expression level that is considered to be CAR positive for the ON and OFF switch, and low CAR expression level in the EXP switch.

The EXP Switch exhibits an increase in mean CAR expression across all recombinase-positive cells, though only approximately 20% of the cells express CAR. Indeed, all three circuits demonstrate that our populations are not homogenously expressing all components of the circuit. In addition to variations in recombinase and CAR expression (**Supplementary Fig. 2**), not all induced ON Switch cells express CAR (**Fig. 2**). Nor do all uninduced OFF and EXP switch cells express CAR (**Fig. 2**). This population heterogeneity is likely due to transduction inefficiency. Indeed, when we integrated these circuits into Jurkat T cells, which generally exhibit greater ease of transduction, we observed cell populations with greater homogeneity (**Supplementary Fig. 3**).

**Figure 3.**
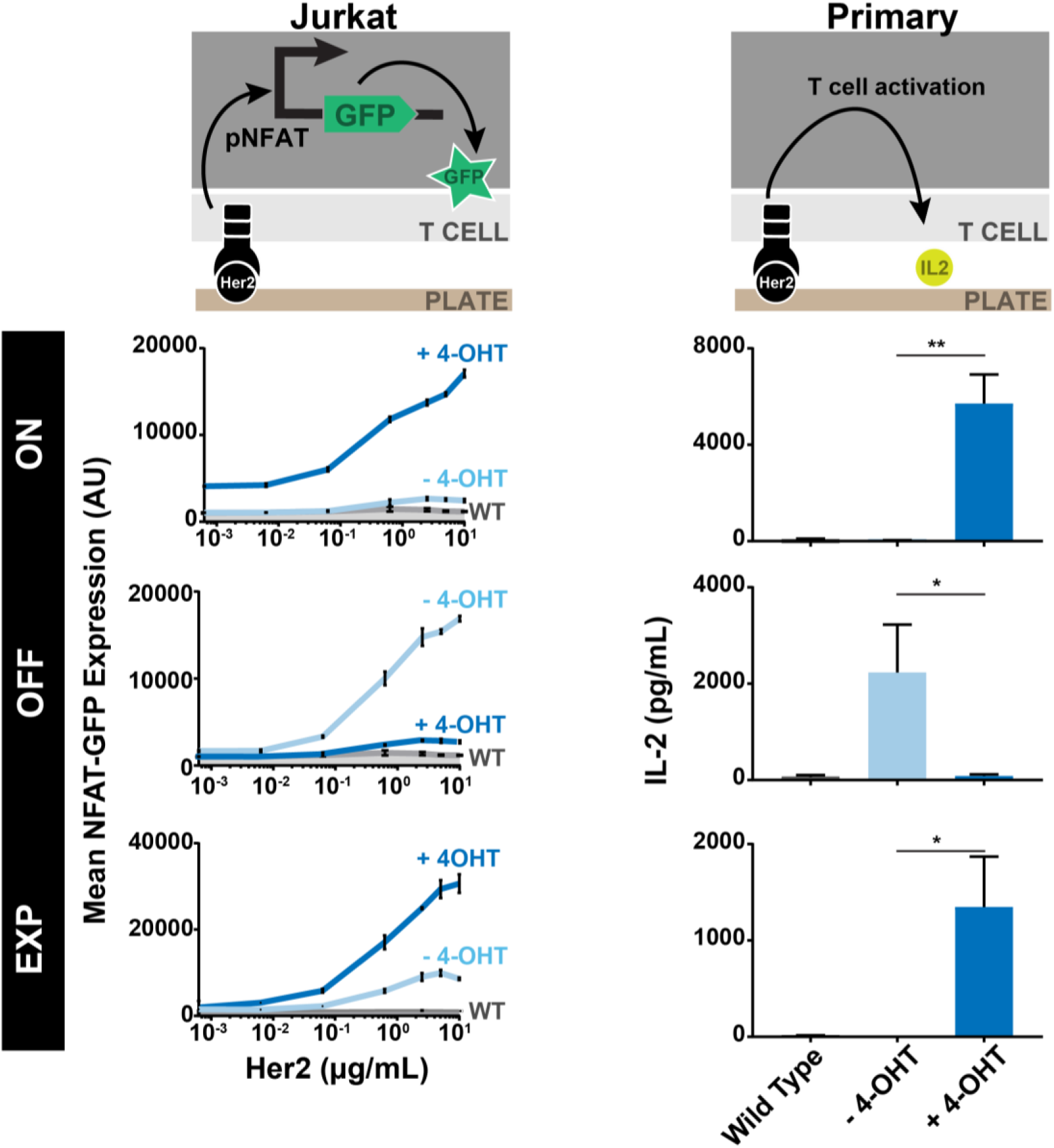
αHer2 CAR activation in T cells containing the recombinase switches. Jurkat T cells (Left Panel) and human primary CD4+ T cells (Right Panel) containing the ON, OFF, or EXP switch controlling expression of an αHer2-CAR were exposed to 4-OHT prior to CAR activation for 5 days (Jurkat, ON and OFF), 8 days (Jurkat, EXP), and 9 days (Primary CD4+, all). Jurkat cells were exposed to different concentrations of plate-bound Her2 protein, and primary T cells were exposed to 5 ug/mL Her2 An NFAT-GFP transcription reporter and IL-2 were measured for Her2 CAR Jurkat and primary T cell respectively. Cells were plated against Her2 antigen in triplicate, and both NFAT-GFP (arbitrary units, AU) and IL-2 (pg/mL) were plotted as mean and standard deviation. Significance in IL-2 production was determined by unpaired, two-tail T-test (* p<0.05, ** p<0.01)

Two key markers of circuit performance are basal switching and switching completeness. Basal switching is best observed in the small population (~8%) of ON Switch that express CAR prior to induction, which indicates a low level basal FlpOER^T2^ activity (**Fig. 2**). This basal level of CAR expression appears to be connected to a higher level of recombinase expression (**Supplementary Fig. 2**). Switching completeness is better observed in the OFF switch, where less than 10% of cells still express CAR at the end of induction. In addition to these markers of circuit activity, we observed high viability and cell growth comparable to wild-type for switch-expressing cells over the course of induction (**Supplementary Fig. 4**), suggesting that genotoxicity of Flp recombinase is minimal.

**Figure 4.**
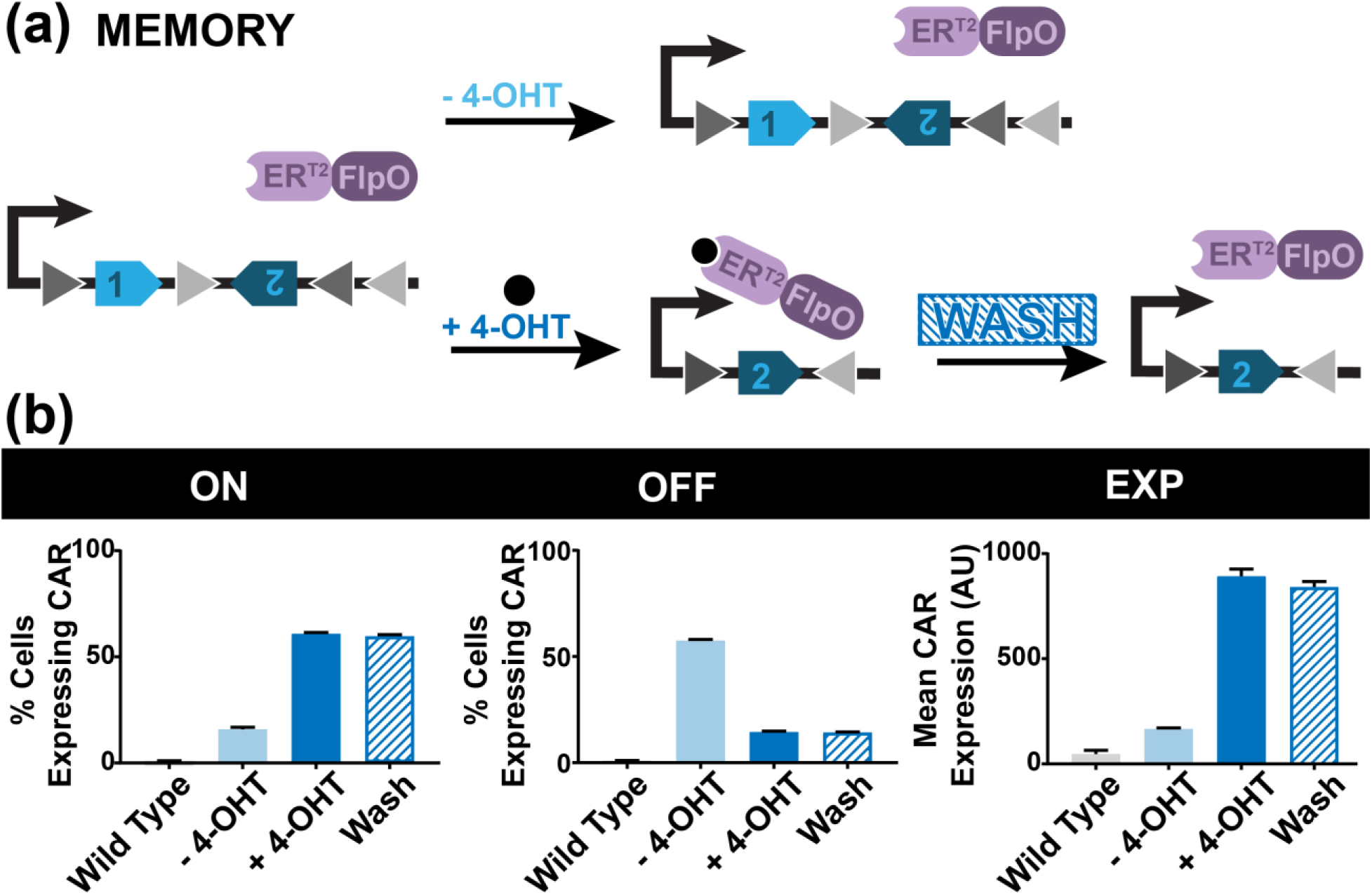
Recombinase switches in primary T cells maintained CAR expression memory after the removal of the inducer. (**a**) Experimental work flow characterizing the switch circuit memory. (**b**) CAR expression from the ON, OFF, or EXP switch with or without washing the 4-OHT after 2 days of 4-OHT exposure. CAR expression was measured 15 days after the initial drug addition. Samples were obtained in triplicate from each induced or non-induced culture and then plotted as mean and standard deviation.

### Changes in CAR expression impact Jurkat NFAT signaling

The induced changes in CAR expression by our switches are expected to alter T cell responses to antigen stimulation. We quantified the activation of circuit-expressing T cells by stimulating them with plate-bound Her2 antigen and measuring the NFAT transcription reporter level in Jurkat T cells (**Fig. 3**). We observed that induced ON switch T cells, which express a CAR after 4-OHT-induction, can be activated with Her2. There was a low level of NFAT reporter expression at higher Her2 doses that suggests some basal CAR expression in uninduced cells, which could be due to a low level of basal recombinase activity. With the OFF switch, we observed that 4-OHT-induced cells have low NFAT reporter activity when stimulated with Her2, corresponding to the loss of CAR expression. The remaining level of NFAT activity in induced cells may be due to incomplete FLEx circuit switching, which could result from factors such as inaccessibility to the integration site. These results are mirrored in ON and OFF switch CD4+ primary T cells that are activated 10 days after induction, where activation is measured via IL-2 production (**Fig. 3**).

For EXP switch-expressing Jurkat T cells, we observed that the increase in CAR expression after 4-OHT induction led to a corresponding increase in NFAT-GFP expression (**Fig. 3**). Interestingly, while we observed this effect for a low-affinity Her2-CAR (C6.5G98A, K_D_ = 3.2×10^−7^)^3^, when the expression level switch is applied to Her2-CARs with increased affinity (C6.5, K_D_ = 1.6×10^−8^; C6MH3-B1, K_D_ = 1.2×10^−10^; C6-B1D2, K_D_ = 1.5×10^−11^)^3^, the effect on NFAT-GFP activation levels is mitigated at high levels of Her2 antigen, illustrating that the level of T cell activation is dually modulated by the CAR expression level and its antigen-scFv affinity (**Supplementary Fig. 5**), corroborating similar observations by others^3, 16, 37, 38^. Primary CD4+ T cells expressing the EXP switch also demonstrate an increase in IL-2 production when activated against Her2. Expression of IL-2 in uninduced EXP cells was low and similar to wild-type, which may be due to a combination of the low CAR expression within the cells, the small population of cells expressing CAR overall, and the low affinity of the CAR.

**Figure 5.**
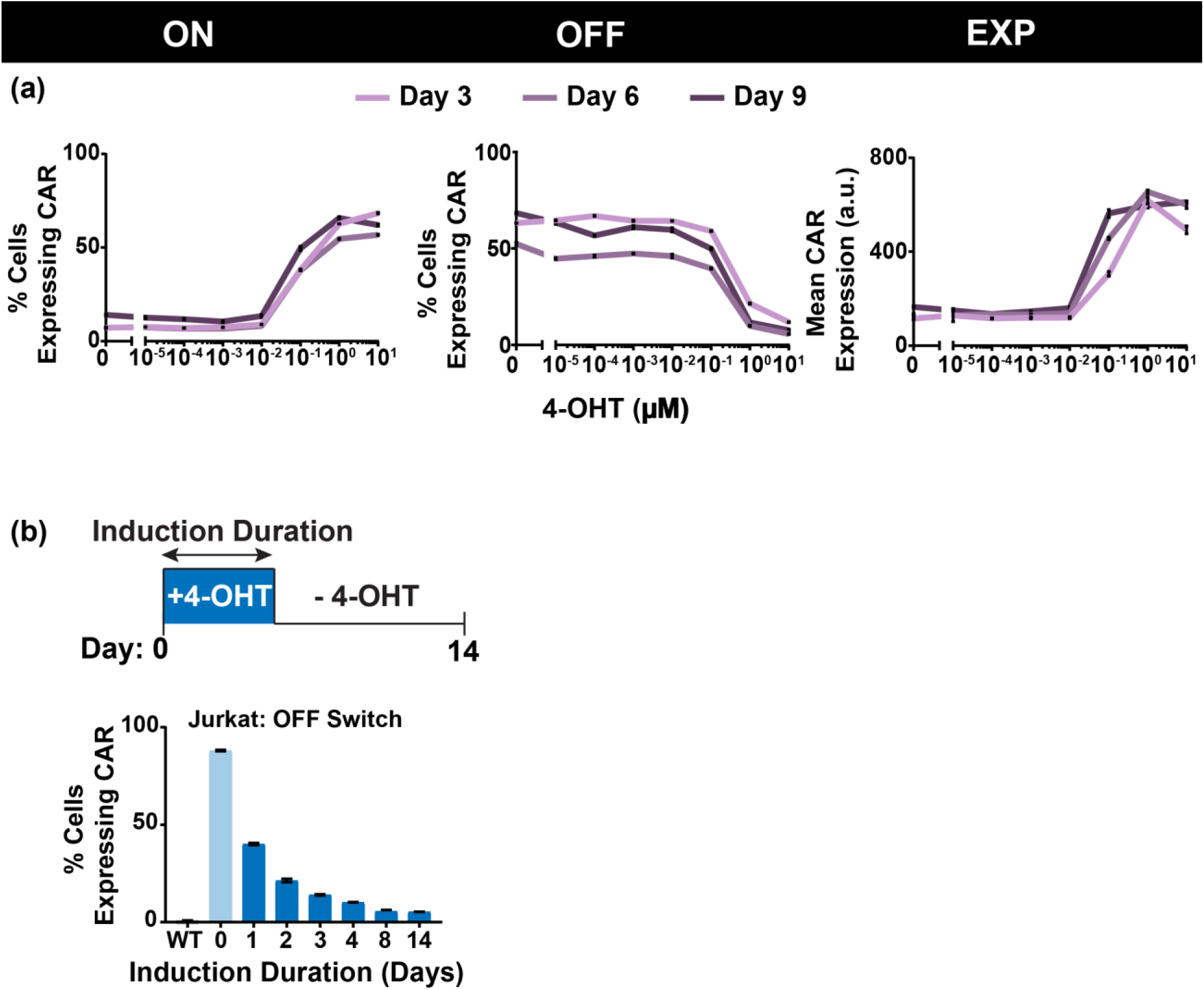
The dose and duration of 4-OHT induction govern the CAR expression from the recombinase switches. (**a**) CAR expression as a function of 4-OHT concentration for the ON, OFF, and EXP switch. Different lines represent the CAR expression at 3, 6 or 9 days after 4- OHT exposure. (**b**) CAR expression as a function of 4-OHT exposure duration. Cells were washed at the indicated day after 4-OHT exposure, and CAR expression was measured 14 days after the initial drug addition. Samples in (a) and (b) were obtained in triplicate from each induced or non-induced culture and then plotted as mean and standard deviation.

### Switch cells exhibit memory when drug is removed

One unique feature of the recombinase-based switch that is not found in other existing gene switches is memory: removal of the drug will maintain any changes made to the cell. Cells that have been switched on will stay on, cells that have been switched off will stay off, and cells that now express greater CAR will continue to express greater CAR. We confirmed this feature for the ON, OFF, and EXP switches in human primary CD4+ T cells. Induced cells that were washed of cells 2 days after induction maintained the same level of CAR expression 15 days post-induction as cells that had been continuously induced for 15 days (**Fig. 4**).

### Tunability of switching can be driven by drug dosage and duration

While these circuits ostensibly provide a two-state system, the percentage of cells that switched states may be tunable by applying the inducer at different concentrations. Therefore, we treated recombinase-positive primary CD4+ T cells with a range of 4-OHT dosages and showed that the percentage of cell population switched states in the ON and OFF switch can indeed be modulated (**Fig. 5a**). In addition, the level of mean CAR expression in the EXP switch can be further tuned via drug dosage.

In cases where the switch takes longer to work, the memory feature enables further tuning of switch dynamics. In particular, varying the **duration** of drug dosage resulted in varying degrees of switching in Jurkat OFF Switch cells (**Fig. 5b, Supplemental Fig. 6**). However, these results did not extend to primary T cells, where all switches expressed the same level of CAR when induced for only 2 days compared to cells that were induced for much longer (**Figure 4**). Jurkat ON and EXP switch cells also did not exhibit drug duration tunability (**Supplemental Fig. 6**).

## Discussion

We have presented a recombinase-based genetic circuit system that allows for increased control of primary T cell behavior. The circuits we have presented here are a lentiviral compatible system with memory capability such that continual addition of the drug is not required for maintenance of any desired changes to the cellular state. This capacity for memory would be particularly important for therapeutic strategies that require permanent changes to be made to the cell, such as permanently shutting off expression of a particular gene, enabling only a temporary drug intake to make changes in lieu of asking a patient to continuously consume the drug. These switches can be used to turn a gene ON or OFF, as well as to stably alter the level of gene expression.

The activity of these circuits provides a number of advantages for patients who require a tunable but permanent change made to the behavior of their engineered T cells. While transiently inducible systems, such as a the drug-dimerizable “ON Switch” CAR, can be powerful forms of control, the memory capacity of the circuits described here will enable the development of a broader class of therapeutic strategies that can make long-lasting changes to T cell behavior without requiring the patient to constantly consume the inducing drug. Minimizing the duration of drug exposure could reduce the complexity of the treatment and the potential toxic side effect of the drug-inducer.

In addition, the tunability of the circuit behavior enables not only the pre-programmed State 1 and State 2 expressed within the circuit, but also a range of CAR expression across the population that could further tune therapeutic activity within the patient. The potential of induction duration to tune circuit behavior may also expand the available therapeutic range. At a 4-OHT concentration of 1 µM, we were only able to observe this duration dosage property in OFF Switch Jurkat T cells. But exploring the combined parameter space of drug dosage (particularly lower drug concentrations) and drug duration, particularly *in vivo*, may reveal more tunable behavior for all three switches in primary T cells. Indeed, *in vivo* use of these switches will require greater study and investigation into the kinetics of both drug delivery and induction of cells.

Our genetic circuits also enable more complex T cell therapeutics by providing a platform to program two different therapeutic states into a cell simply by expressing a particular gene of choice within the switch cassette. This increased complexity will expand therapeutic strategies that can be implemented, as it will enable easy bedside switching between two states of therapy based on a patient’s individual needs. While we have focused on its use with CARs in this work, our circuit is also compatible with other forms of T cell therapies that could benefit from control of other genes.

Our use of FlpOER^T2^ takes advantage of the power of drug-inducible recombinases to create powerful genetic technologies, as well as the improved viability of cells expressing FlpO over the commonly used Cre recombinase. However, one potential limitation of recombinases is their immunogenicity due to their non-human origins (FlpO, for example, is derived from yeast). To mitigate transgene immunogenicity, one strategy is to leverage genome editing tools to eliminate the gene B2M, an important component for antigen display through class I human leukocyte antigen (HLA)^39, 40^. This strategy has also been combined with HLA-E overexpression to reduce immunogenicity against pluripotent stem cells^41^. This approach would not only enable the development of “universal T cells” and provide a bank of “off-the-shelf” therapeutic T cells, it would enable the incorporation of genetic technologies comprised of proteins from diverse organisms. Therefore, with the rapid advancement in genome editing technologies, we are confident that transgene immunogenicity derived from FlpO can be addressed.

Optimized use of our recombinase-based switching will rely on careful consideration of the potential application. In particular, there are situations where low levels of basal activity or incomplete switching may carry enough risk that this approach will not be applicable. Careful tuning of recombinase expression or activity may provide further avenues to address these limitations, as could the incorporation of inducible kill switches to selectively kill cells that are not behaving as directed.

T cell therapies will require us to not just rely on the mechanics of the immune system, but to understand the intricacies that are available and necessary for us to fine-tune in order to create a safe and effective treatment. With many developments and tools focused around developing one facet of control, having a platform of genetic circuits that can be applied in different ways creates a wider array of options available to implement T cell therapies.

## Methods

### Circuit Construction

The circuit we described is comprised of two parts: the inducible recombinase and the FLEx switch. These components were cloned into separate lentiviral backbones using a combination of Gibson and traditional molecular cloning methods. The FlpOER^T2^ recombinase was cloned into the backbone followed by a T2A ribosomal skip sequence and an mTAG-BFP fluorescent marker.

The FLEx switch was designed using the *frt* and *f3* recombination sites. For the ON and OFF switches, the chimeric antigen receptor sequence was inserted between the recombination sites. The SFFV promoter was used to drive FLEx switch (and thus, CAR) expression. For the expression level switch, the EF1α promoter was inserted between the recombination sites such that the reverse promoter orientation was encoded in the 5’ to 3’ direction. The CAR was expressed downstream of the FLEx/reverse EF1α promoter.

### Primary T cell isolation and transduction

Blood was obtained from the Boston Children’s Hospital, and primary CD4+ T cells were harvested using the STEMCELL CD4+ enriched cocktail in conjunction with the RosetteSep system. T cells were preserved at −80°C in 90% FBS (Gibco) and 10% DMSO. T cells were maintained X-Vivo 15 media (Lonza) supplemented with 5% Human AB Serum (Valley Biomedical), 10 mM N-acetyl L-Cyteine (Sigma), and 55 µM 2-mercaptoethanol (Gibco). Through thawing and transduction, T cells were maintained with 100 units/mL recombinant IL-2 (Tecin, NCI BRB Preclinical Repository) and then 50 units/mL post-transduction.

Human embryonic kidney (HEK) 293 FT cells were transfected with lentiviral packaging plasmids and either the FLEx switch plasmid or the inducible recombinase plasmid in a T175 flask using polyethylenimine (PEI). One day after transfection, media was replaced with Ultraculture media (Lonza) supplemented with 100 U/mL Penicillin+100 µg/mLStreptomycin (Corning), 2mM L-Glutamine (Corning), 50 mM Sodium Butyrate (Alfa Aesar), and 1 mM Sodium Pyruvate (Lonza), and virus was collected three and four days after transfection by collecting and spinning the media, and retaining the supernatant. Virus was concentrated through ultracentrifugation with 20% sucrose (Sigma) for two hours at 4 °C and 22,000xg.

T cells were thawed two days prior to transduction and activated with CD3/CD28 Dynabeads (Gibco) one day prior. Cells were transduced via spinfection: using half of concentrated virus, both inducible recombinase and switch viruses were spun onto the well of a 6-well retronectin (Clontech)-coated plates for 90 minutes at 1200xg. Activated primary T cells were then spun onto the virus plates for 60 minutes at 1,200xg.

### Jurkat T cell maintenance and transduction

Through transduction and general maintenance, Jurkat T cells were maintained in RPMI media (Lonza) supplemented with 5% fetal bovine serum (Gibco), 2mM glutamine, and 100 U/mL penicillin+100 µg/mL streptomycin. Through analysis, cells were maintained in RPMI media supplemented with 10% fetal bovine serum and 2mM glutamine.

Lentiviral transduction was used to produce T cell lines containing full circuitry (inducible recombinase and designated ON/OFF/EXP switch). HEK293FT cells were transfected via PEI in a 6 well plate with lentiviral packaging plasmids and the circuit component lentiviral plasmid to produce virus containing the specified component. Virus was collected three days after transfection.

Approximately 500,000 Jurkat NFAT cells—a line produced by the Weiss lab at UCSF^42^ to express an NFAT-GFP activation reporter—were infected with 500 µL of the recombinase virus and 500 µL of the switch for co-transduction of the entire circuit. Transduced Jurkat-NFAT cell were diluted with media one day after infection and then collected three days after infection.

### Switch Induction with 4-OHT

Cells were induced with 4-hydroxytamoxifen (4-OHT, Sigma), a metabolite of tamoxifen, in methanol solution. All induction experiments were conducted with 1 µM 4-OHT except for dose response experiments, which were conducted with a 4-OHT concentration range from 10^−5^ to 10 µM. For induction time courses, cells were induced at a starting concentration of 200,000 cells/mL and maintained between 200,000-1,200,000 cells/mL with media containing 1 µM 4- OHT. Uninduced cells were also plated and maintained at the same concentrations in inducer-negative media.

To test effect of induction duration, cells were induced at a starting concentration of 800,000 cells/mL. For each day up until 4 days post-induction, a fraction of induced cells were removed. Removed cells were then spun down, washed with 5 mL inducer-negative media, and then resuspended to a concentration of 800,000 cells/mL in inducer-negative media. Induced cells were then diluted 1:2. On day 4, all cells were diluted 1:8, and 8 days post-induction, one last batch of induced cells were washed. Through the rest of the experiment, cells were maintained between 200,000-1,600,000 cells/mL.

### CAR Activation with plate-bound Her2 protein

Target antigen was plated on 96 well, tissue culture-treated flat bottom plates in Dulbecco’s phosphate buffered saline (PBS, Corning) for two hours at 37°C. Wells were then washed two times with PBS, and 200,000 cells at a concentration of 1,000,000 cells/mL were plated overnight. To remove supplemental IL-2 prior to plating, primary CD4+ T cells were washed two times and ultimately plated with IL-2-negative media. Cells were plated with or without 4-OHT in accordance with their induction conditions.

### Flow cytometry sorting and analysis

Cells were sorted for BFP-positive expression using the SH800 Cell Sorter (Sony. BFP-FL1 channel). To measure circuit switching dynamics and NFAT-GFP expression were measured via flow cytometry (Attune NxT Flow Cytometer, Thermo Fisher Scientific. BFP-VL1 channel, PE-YL1 channel, mCherry-YL2 channel, GFP-BL1 channel). Results were analyzed using FlowJo 10.0.7 (FlowJo, LLC). CAR expression in the EXPression Level switch was characterized via an mCherry tag directly connected to the CAR. CARs in ON and OFF switches did not contain the mCherry tag, and expression was measured by staining for a myc epitope tag expressed in the extracellular portion of the CAR. Staining was done using a PE-conjugated human c-myc antibody (R&D Systems IC3696P) at a concentration of 10 μL antibody/10^6^ cells.

### IL-2 ELISA

IL-2 production by CD4+ primary T cells was measured using an ELISA kit (BD 550611) according to manufacturer’s instruction. 100 μL of supernatant from each sample was collected 19 hours post-induction and frozen in −80°C prior to quantification with the ELISA.

